## Supplementary material for "A simple thermodynamic description of phase separation of Nup98 FG domains": Supp.

Contents:

**Supplementary Figures 1-5** with legends
**Supplementary Tables 1-5**
**References**

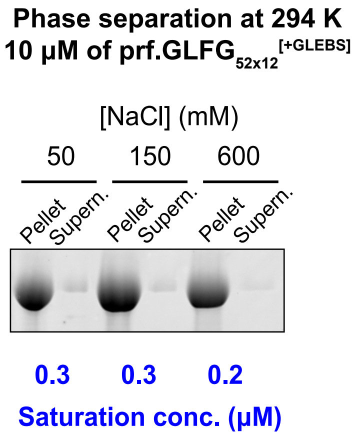


**Supplementary Figure 1: prf.GLFG$_{52x12}$$^{[+GLEBS]}$ is a host phase with low saturation concentration.** 10 µM dilutions of prf.GLFG$_{52x12}$$^{[+GLEBS]}$ were prepared in buffers containing the indicated concentration of NaCl and centrifuged at the same temperature (21°C/294 K). SDS samples of the obtained pellets (FG phase) and supernatants (soluble content) were loaded for SDS-PAGE at equal ratio (7%), followed by Coomassie blue staining. Saturation concentrations for each condition were determined as described in the main text, and are shown in blue. A full scan of the gel with molecular weight markers is provided in the Source Data file. Experiments for each condition were repeated two times on independent samples with similar results and the mean values are shown.

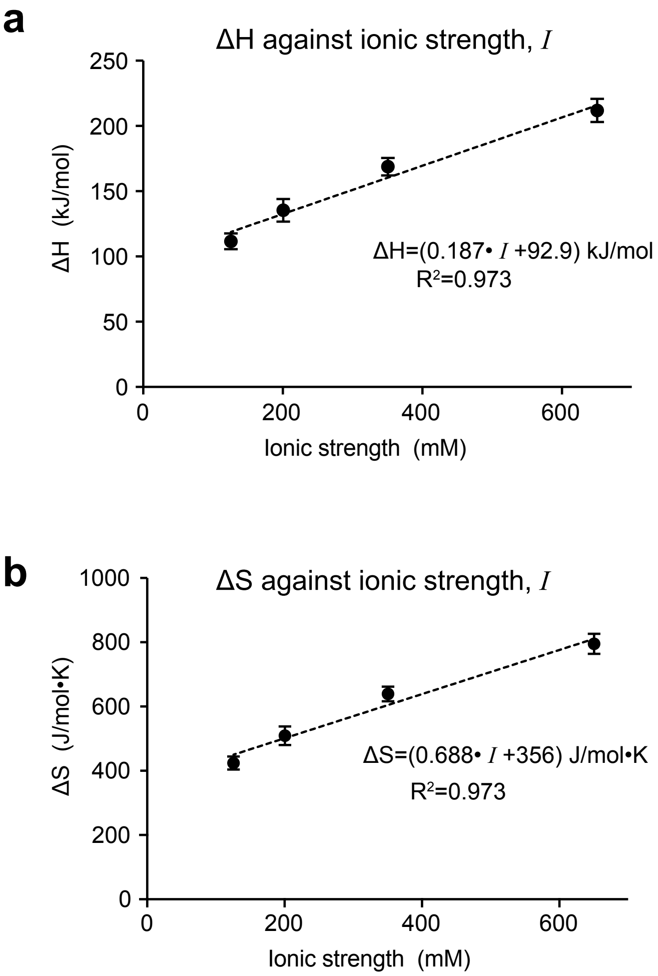


**Supplementary Figure 2: *ΔH* and *ΔS* of phase separation of prf.GLFG$_{52x12}$ against ionic strength.** *ΔH* and *ΔS* corresponding to different salt concentrations were obtained by plotting ln ($C_{sat}$ / $C_{dense}$) against $1/T$ (see Fig. 4a and Supp. Table 2). Here the *ΔH* **(a)** and *ΔS* **(b)** (data are presented as best-fit values with standard errors (S.E.) of fitting as the error bars) are plotted against the total ionic strength ($I$) of the assay buffer (=contribution from 20 mM NaPi + contribution from NaCl), where:

$$I = \frac{1}{2}\sum_{i=1}^{n} c_i z_i^2$$

$c_i$ is the molar concentration of ion $i$ (among $n$) and $z_i$ the corresponding charge.

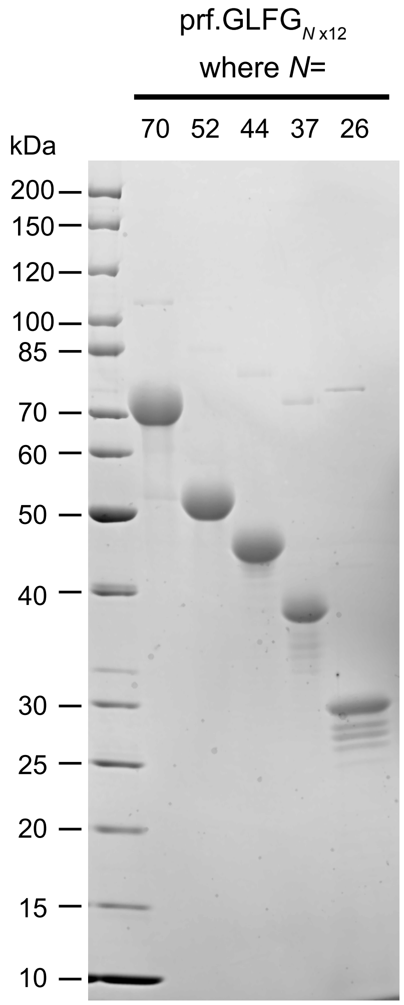


**Supplementary Figure 3: FG domain variants of perfect repeats on a Coomassie-stained SDS-gel.** Different FG domain variants were constructed, each composed of a given number ($N$) of connected perfect repeats of a 12mer peptide: <GGLFGGNTQPAT>. An SDS gel stained by Coomassie blue shows each variant before phase separation. 6 µg of prf.GLFG$_{26\text{x}12}$ was loaded and 4 µg was loaded for the rest. Expected molecular masses are: 77.4, 57.5, 48.9, 41.2 and 29.1 kDa for $N$ = 70, 52, 44, 37 and 26 respectively. Note that for these FG domain variants, the apparent sizes estimated from the gel are lower than the expected molecular weights, due to their unusual amino acid compositions.

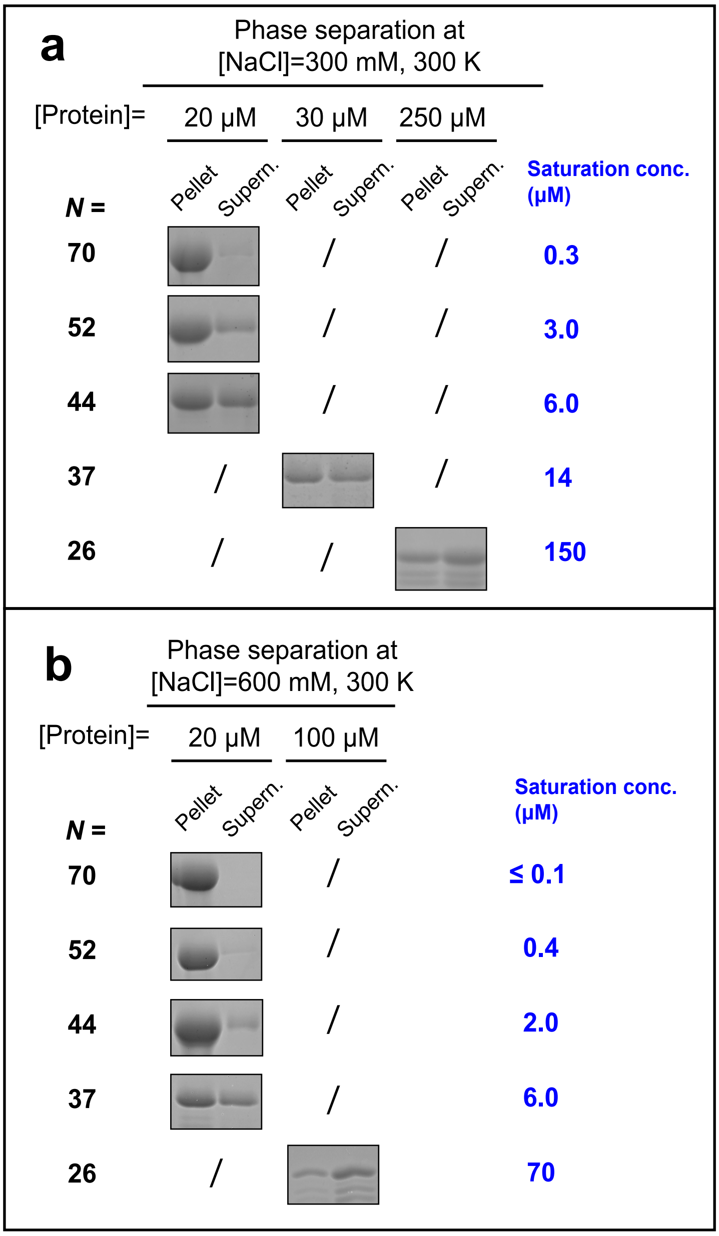


**Supplementary Figure 4: Measurements of saturation concentrations at 300 and 600 mM NaCl.** Variants containing different number of perfect repeats (*N*) were centrifuged at 300 mM **(a)** and 600 mM NaCl **(b)**. SDS samples of the obtained pellets (FG phase) and supernatants were loaded for SDS-PAGE at equal ratio, followed by Coomassie blue-staining. Values of saturation concentration for each condition were determined as described in the main text, and are shown in blue. Assay concentrations of the variants are as indicated. Marked with "/": not determined. Assay temperature was 300 K for both **(a)** and **(b)**. Full scans of gels with molecular weight markers are provided in the Source Data file. Experiments for each condition were repeated two times on independent samples with similar results and the mean values of saturation concentration are shown. The results are plotted in Fig. 7a.

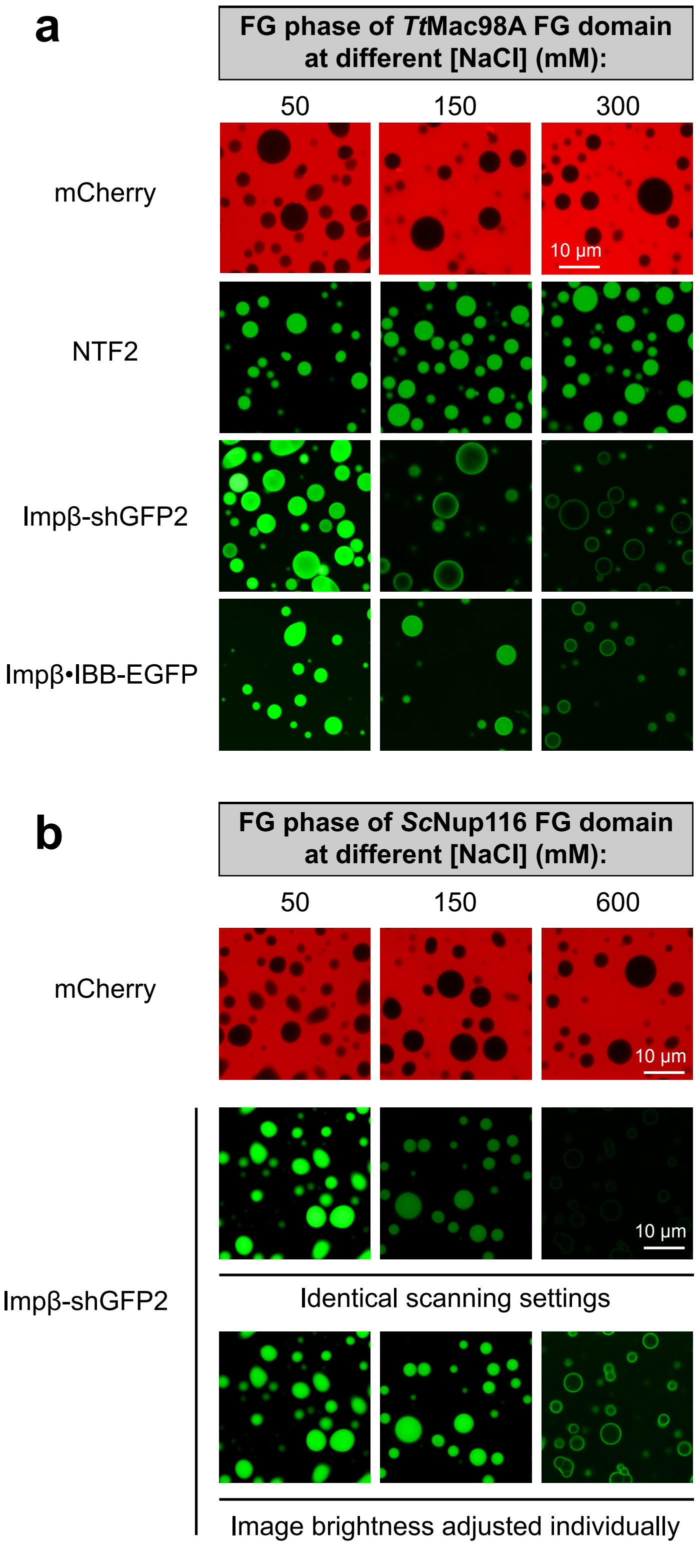


**Supplementary Figure 5: Permeation selectivity of FG phases at different [NaCl].**
FG phases assembled by wildtype FG domains of *Tt*Mac98A **(a)** and *Sc*Nup116 **(b)** were challenged with the indicated permeation probes at different salt concentrations. Impβ·IBB-EGFP: a 130 kDa complex of Impβ and a standard EGFP containing an IBB domain for Impβ-binding. Identical scanning settings and image processing were applied for the same permeation probe. One more image set is included for Impβ-shGFP2 in **(b)** (as indicated) whereas the image brightness was adjusted between the set to show the weaker signals.

| Symbol | Meaning |
|---|---|
| $a$ | $d(\Delta H)/d(C_{NaCl})$, slope of a plot of $\Delta H$ against $C_{NaCl}$ (Fig. 4b),<br>enthalpy change per unit NaCl in solvent (Eqs.5,7 &8) |
| $b$ | intercept of a plot of $\Delta H$ against $C_{NaCl}$ (Fig. 4b),<br>enthalpy when $C_{NaCl}$ =0 (Eqs.5,7 &8) |
| $c$ | $d(\Delta S)/d(C_{NaCl})$, slope of a plot of $\Delta S$ against $C_{NaCl}$ (Fig. 4c),<br>entropy change per unit NaCl in solvent (Eqs.6,7 &8) |
| $d$ | intercept of a plot of $\Delta S$ against $C_{NaCl}$ (Fig. 4c),<br>entropy when $C_{NaCl}$ =0 (Eqs.6,7 &8) |
| $C_{dense}$ | concentration of the condensed FG phase expressed in molar concentration of the FG domain molecules |
| $C_{sat}$ | saturation concentration of phase separation (expressed in molar concentration), *aka* threshold concentration |
| $C_{NaCl}$ | Molar concentration of NaCl (=[NaCl]) in an aqueous buffer |
| *CMC* | critical micelle concentration expressed in molar concentration |
| $K$ | distribution constant |
| $k_B$ | the Boltzmann constant |
| $N$ | Number of connected repeats in a perfectly repetitive FG domain |
| $p$ | $p$-value |
| $P$ | partition coefficient |
| $R$ | the gas constant |
| $R^2$ | R-squared value (coefficient of determination) |
| $T$ | phase transition temperature, *aka* cloud point temperature |
| $w$ | $d(\Delta H)/dN$, slope of a plot of $\Delta H$ against $N$ (Fig. 6b),<br>enthalpy change per repeat unit (Eqs.9,11,12) |
| $x$ | a constant representing the intercept of a plot of $\Delta H$ against $N$ (Fig. 6b) (Eqs.9 &12) |
| $y$ | $d(\Delta S)/dN$, slope of a plot of $\Delta S$ against $N$ (Fig. 6c),<br>entropy change per repeat unit (Eqs.10,11,12) |
| $z$ | a constant representing the intercept of a plot of $\Delta S$ against $N$ (Fig. 6c) (Eqs.10 &12) |
| $\Delta G$ | Gibbs free energy change for phase separation of a given FG domain |
| $\Delta G_{part.}$ | Gibbs free energy change for partition into an FG phase |
| $\Delta G_{repeat}$ | Energy contributed by one repeat unit to Gibbs free energy change for phase separation/ partition |
| $\Delta H$ | enthalpy change for phase separation |
| $\Delta S$ | entropy change for phase separation |

**Supplementary Table 1: Symbols used in this manuscript.**

| $C_{NaCl}$ (mM) | Slope ± S.E. ($10^3$·K) | $\Delta H$ ± S.E. (kJ/mol) | Intercept ± S.E. (Dimensionless) | $\Delta S$ ± S.E. (J/mol·K) |
|---|---|---|---|---|
| 75 | 13.2 ± 0.7 | 110 ± 6 | -50.1 ± 2.4 | 416 ± 20 |
| 150 | 15.9 ± 1.0 | 132 ± 8 | -60.2 ± 3.5 | 501 ± 29 |
| 300 | 20.2 ± 0.8 | 168 ± 7 | -75.9 ± 2.8 | 631 ± 23 |
| 600 | 25.3 ± 1.1 | 210 ± 9 | -94.7 ± 3.7 | 788 ± 31 |

**Supplementary Table 2: Thermodynamic parameters for phase separation of prf.GLFG$_{52x12}$.** Parameters were obtained by plotting ln $(C_{sat}/\ C_{dense})$ against $1/T$ (van't Hoff plots shown in Fig. 4a, based on a DLS dataset):

$$\Delta H = slope \times R$$

$$\Delta S = -intercept \times R$$

Data are presented as best-fit values ± standard errors (S.E.) of fitting. Note that the S.E. are typically less than 6% of the best-fit values. Eqs. 5 and 6 in the main text represent the trends for $\Delta H$ against $C_{NaCl}$ and $\Delta S$ against $C_{NaCl}$ respectively.

| Number of repeats, *N* | *P* | $C_{dense}$/ $C_{sat}$ |
|---|---|---|
| 70 | 2400 | 1300 |
| 52 | 600 | 380 |
| 44 | 300 | 180 |
| 37 | 80 | 90 |
| 26 | 30 | Not measurable |
| 18 | 11 | Not measurable |
| 13 | 6 | Not measurable |
| 7 | 4 | Not measurable |

**Supplementary Table 3: Comparisons of orthogonal datasets.**
Results obtained from two orthogonal methods are listed and compared. In the first method, partition coefficients (*P*) of fluorescently labelled variants, each with a given number of FG repeats (*N*), into a pre-formed "host" FG phase, assembled by prf.GLFG$_{52x12}^{[+GLEBS]}$ were measured (see Fig. 1). In the second method, the phase separation of variants with $N$ = 70, 52, 44, and 37 was analysed individually (Fig. 5) and for each, $C_{sat}$ was measured to calculate $C_{dense}$ : $C_{sat}$.
For comparison of the two orthogonal datasets, the values listed here were obtained under the same salt concentration (150 mM NaCl) and temperature (294 K).

| Number of repeats, $N$ | $C_{NaCl}$ (mM) | Slope ± S.E. ($10^3$·K) | $\Delta H$ ± S.E. (kJ/mol) | Intercept ± S.E. (Dimensionless) | $\Delta S$ ± S.E. (J/mol·K) |
|---|---|---|---|---|---|
| 70 | 150 | 14.5 ± 0.7 | 121 ± 6 | -56.5 ± 2.4 | 470 ± 20 |
| 52 | | 11.9 ± 0.4 | 99 ± 4 | -46.3 ± 1.5 | 385 ± 12 |
| 44 | | 10.1 ± 0.6 | 84 ± 5 | -39.7 ± 2.1 | 330 ± 18 |
| 37 | | 9.0 ± 0.4 | 76 ± 3 | -35.4 ± 1.2 | 295 ± 10 |
| 26 | | 6.7* | 56* | -26.4* | 220* |

**Supplementary Table 4: Thermodynamic parameters for phase separation of perfectly repetitive variants.** Parameters were derived from van't Hoff plots shown in Fig. 6a. which are based on a centrifugation dataset (Fig. 5) with different FG domain variants each composed of a given number ($N$) of connected perfect repeats of a 12mer peptide: <GGLFGGNTQPAT>. Calculations are as described in the main text. Data are presented as best-fit values ± standard errors (S.E.) of fitting. *S.E. of parameters for the 26x repeat cannot be determined due to insufficient data points. Eqs.9 and 10 in the main text represent the trends for $\Delta H$ against $N$ and $\Delta S$ against $N$ respectively.

| Protein name | Plasmid | Encoding for | Used in figures | Reference |
|---|---|---|---|---|
| prf.GLFG$_{52x12}$* | pSNG064 | His$_{14}$-ZZ-*Sc*SUMO-prf.GLFG$_{52x12}$ | 3,4,5,6,7 S2,S3,S4 | 1 |
| prf.GLFG$_{52x12}$-Cys* | pSNG102 | His$_{14}$-ZZ-*Sc*SUMO-prf.GLFG$_{52x12}$-Cys | 1,2 | 1 |
| prf.GLFG$_{70x12}$-Cys* | pSNG111 | His$_{14}$-ZZ-*Sc*SUMO-prf.GLFG$_{70x12}$-Cys | 1,5,6,7, S3,S4 | this study |
| prf.GLFG$_{44x12}$-Cys* | pSNG105 | His$_{14}$-ZZ-*Sc*SUMO-prf.GLFG$_{44x12}$-Cys | 1,2,5,6,7, S3,S4 | this study |
| prf.GLFG$_{37x12}$-Cys* | pSNG106 | His$_{14}$-ZZ-*Sc*SUMO-prf.GLFG$_{37x12}$-Cys | 1,2,5,6,7, S3,S4 | this study |
| prf.GLFG$_{26x12}$-Cys* | pSNG107 | His$_{14}$-ZZ-*Sc*SUMO-prf.GLFG$_{26x12}$-Cys | 1,2,5,6,7, S3,S4 | this study |
| prf.GLFG$_{18x12}$-Cys* | pSNG108 | His$_{14}$-ZZ-*Sc*SUMO-prf.GLFG$_{18x12}$-Cys | 1,2 | this study |
| prf.GLFG$_{13x12}$-Cys* | pSNG109 | His$_{14}$-ZZ-*Sc*SUMO-prf.GLFG$_{13x12}$-Cys | 1,2 | this study |
| prf.GLFG$_{7x12}$-Cys* | pSNG110 | His$_{14}$-ZZ-*Sc*SUMO-prf.GLFG$_{7x12}$-Cys | 1,2 | 1 |
| His$_{18}$-prf.GLFG$_{52x12}$ [+GLEBS] | pSNG038 | His$_{18}$- prf.GLFG$_{52x12}$[+GLEBS]-Cys | 1,8,S1 | 2 |
| His$_{18}$-*Sc*Nup116 FG domain | pHBS698 | His$_{18}$-ScNup116$_{1\text{-}736}$-Cys | 2,S5 | 3 |
| His$_{18}$-*Tt*MacNup98A (Mac98A) FG domain | pHBS418 | His$_{18}$-*Tt*MacNup98A$_{1\text{-}666}$-Cys | 2,3,S5 | 3 |
| His$_{18}$-*Xt*Nup98 FG domain | pHBS553 | His$_{18}$-*Xt*Nup98$_{1\text{-}485}$-Cys | 2,3 | this study |
| His$_{18}$-*Bf*Nup98 FG domain | pHBS505 | His$_{18}$-*Bf*Nup98$_{1\text{-}478}$-Cys | 2,3 | 3 |
| His$_{14}$-*At*Nup98B FG domain | pHBS383 | His$_{14}$-TEV-*At*Nup98B$_{1\text{-}668}$-Cys | 2,3 | 3 |
| His$_{14}$-*Tb*Nup158 FG domain | pHBS249 | His$_{14}$-TEV-*Tb*Nup158$_{1\text{-}565}$-Cys | 2 | 3 |
| mCherry* | pSF779 | His$_{14}$-TEV-mCherry-Cys | 8,S5 | 3 |
| rat (*Rattus norvegicus*) NTF2 | pDG2121 | *Rn*NTF2 | 8,S5 | 4 |
| shGFP2-Impβ * | pSF2051 | His$_{14}$-*Bd*SUMO-shGFP2-*Sc*Kap95 | 8,S5 | this study |
| GFP$^{NTR}$_3B7C* | pDG2779 | His$_{14}$-*Bd*SUMO-GFP$^{NTR}$3B7C | 8 | 4 |
| Importin β* | pDG2305 | His$_{14}$-MBP-*Bd*SUMO-*Hs*Imp_beta | 8,S5 | 4 |
| IBB-sffrGFP7* | pDG2899 | His$_{14}$-*Bd*SUMO-*Hs*IBB-sffrGFP7 | 8 | 4 |
| IBB-EGFP* | pDG2895 | His$_{14}$-*Bd*SUMO-*Hs*IBB-EGFP | S5 | 4 |

**Supplementary Table 5: Proteins and corresponding bacterial expression constructs used in this study.** Plasmid numbers are unique identifiers. * indicates that a histidine-tag-cleaved (by SUMO or TEV) version of the protein was used.
